# The evolutionary history of plastid outer envelope proteins – a structure-sequence comparison

**DOI:** 10.1101/2025.06.18.660430

**Authors:** Deren Büyüktaş, Jonas A. Mehl, Timo Engelsdorf, Lars M. Voll, Hans-Henning Kunz, Serena Schwenkert, Sophie de Vries

**Affiliations:** Department of Applied Bioinformatics, Institute of Microbiology and Genetics, University of Goettingen, Goldschmidtstraße 1, 37077 Goettingen, Germany; Philipps-Universität Marburg, Department Biology, Molecular Plant Physiology, Karl-von-Frisch-Strasse 8, D-35043 Marburg, Germany; Center for Synthetic Microbiology (SYNMIKRO), Philipps-Universität Marburg, Karl-von-Frisch-Strasse 14, D-35043 Marburg, Germany; Plant Biochemistry, Biology, LMU Munich, Großhadernerstr. 2-4, 82152 Planegg-Martinsried, Germany; Plant Molecular Biology, LMU Munich, Großhaderner Str. 2-4, 82152 Planegg-Martinsried, Germany

## Abstract

Plant metabolism heavily relies on chloroplasts, derived from once free-living organisms. However, how two distinct organisms merged into one remains currently only partially understood. Protein-mediated metabolite exchange across the double-membrane chloroplast envelope is essential for plant cell function. Here, we investigate the evolutionary origins of outer envelope proteins (OEPs) involved in these transport processes. The mosaic nature of the nuclear genome and the deep evolutionary distance since plastid acquisition are major challenges. To address them, we combine sequence-based analyses with emerging structure-based tools, which together enable more sensitive evolutionary comparisons than traditional methods alone. To uncover distinct evolutionary trajectories, we focused on five OEP families: four β-barrel proteins involved in metabolite transport and JASSY, the first published OPDA transporter. We found that the β-barrel proteins were recruited in a stepwise manner and structural homologs of some OEPs point to early recruitment via endosymbiotic gene transfer (EGT). Notably, OEP40 shows a recent structural rearrangement, lacking clear structural homologs in plants other than Arabidopsis, yet retaining sequence conservation across all major land-plant lineages. The JASSY-like family is found across plastid-bearing species, while true JASSY orthologs emerged in embryophytes likely via a stable recruitment of the characteristic lipid-binding domain. Overall, our findings highlight the dynamic nature of the chloroplast outer envelope and show how new functions evolved through structural reshaping and novel domain recruitment. Structure-based approaches thus powerfully complement sequence data, offering more in-depth insight into the evolution of plastid transport systems.

## Introduction

Chloroplasts are vital for plant cells. They produce the carbon building blocks for almost all aspects of plant physiology. All plastids can be traced to a single endosymbiotic event between a heterotrophic, eukaryotic host and a photoautotrophic cyanobacterium. Successful establishment of this relationship was accompanied by massive EGT from the cyanobacterial genome to the nucleus (Timmis et al., 2004). In addition, horizontal gene transfer (HGT) from other prokaryotic sources into the endosymbiont genome and from there to the host is observed (Dagan et al. 2013, Ku et al. 2015, Ponce-Toledo et al. 2019). This complicates tracing the evolutionary origin of genes relevant for chloroplast function. The genetic exchange between the two partners resulted in a mosaic metabolism, which made the two organisms interdependent (Fernie et al. 2024) e.g., chloroplast function relies on nutrients from the cytosol in turn plastids provide fixed carbon and special metabolites. To ensure flawless interaction, plastid outer and inner envelope (OE and IE) membranes harbor a variety of integral proteins assigned to metabolite, ion, and protein transport (Kunz et al. 2024, John et al. 2025) Weber et al. 2005, Bölter 2018).

The abundance of β-barrel proteins in the OE may be especially informative to understand how the cyanobacterial outer membrane was reorganized to allow permanent maintenance of the endosymbiont. In contrast to mitochondria, which mainly rely on the voltage-dependent anion channel (VDAC) as metabolite channel, OEPs appear to have diversified to a greater extent in the course of evolution (Barth et al. 2022, Schwenkert et al. 2023). Despite decades of research, a detailed understanding of the evolutionary recruitments of molecular mechanisms that allowed and shaped the emergence of the plastid remains scarce (Barth et al. 2022). This information must be engraved in the sequence and structure of key proteins. However, strong evolutionary divergence since the emergence of plastids hampers the investigation of its evolutionary history based on sequence data alone. Homology searches using basic local alignment search tools (BLAST) poorly detect ancient relationships where AA sequence similarity has decayed below 30% (Rost, 1999), often referred to as homologs from the “twilight zone”. Novel approaches capitalize on the fact that, constrained by their biological function, protein structures are ≤10-times more conserved than sequence (Illegård et al. 2009, Ruperti et al. 2023). The emerging abundance of predicted protein structures and the success of structure-based method in homology search and sequence alignment (Chen et al. 2025) has renewed the interest of researchers in using the structural information of proteins to directly improve phylogenetic inference (Moi et al. 2023, Baltzis et al. 2025, Bou Dagher et al. 2025). Yet, structural phylogenetics has not been applied towards understanding the evolution of plant cells. We reason that integration of structural and sequence analyses can overcome the low sequence conservation of OEPs and gain new insights into the evolution of the chloroplast OE as well as the origin and recruitment of key proteins necessary for the processes of organellar integration.

## Results and discussion

### The evolutionary history of OEP protein sequences

To test our hypothesis and identify potential pitfalls in structure-based evolutionary analyses, we carried out a comparative proof-of-concept study employing traditional sequence-based phylogenetics and structural analysis in the phylogeny of the green lineage and cyanobacteria. As candidates we selected five OEP families (four β-barrel proteins and one with mixed structure) with specific functions in metabolite transport (Figure 1a): (i) OEP21, the structure of which was recently solved, is selective for triose phosphates and regulated by ATP, likely preventing the loss of metabolites during oxidative stress (Günsel et al. 2023), (ii) OEP24 and (iii) OEP37 exhibit broader substrate specificity, functioning as general solute channels that can partially substitute for VDAC in yeast systems (Röhl et al. 1999, Götze et al. 2006), (iv) OEP40, a candidate transporter for glucose and phosphorylated derivatives, which contributes to major carbohydrate metabolism and the regulation of flowering under cold stress (Harsmann et al. 2016) and (v) JASSY, which has a mixed alpha helical and β-sheets structure, and was recently characterized to shuttle 12-oxo phytodienoic acid (OPDA) (Guan et al. 2019, Schwenkert et al. 2023).

**Figure 1:**
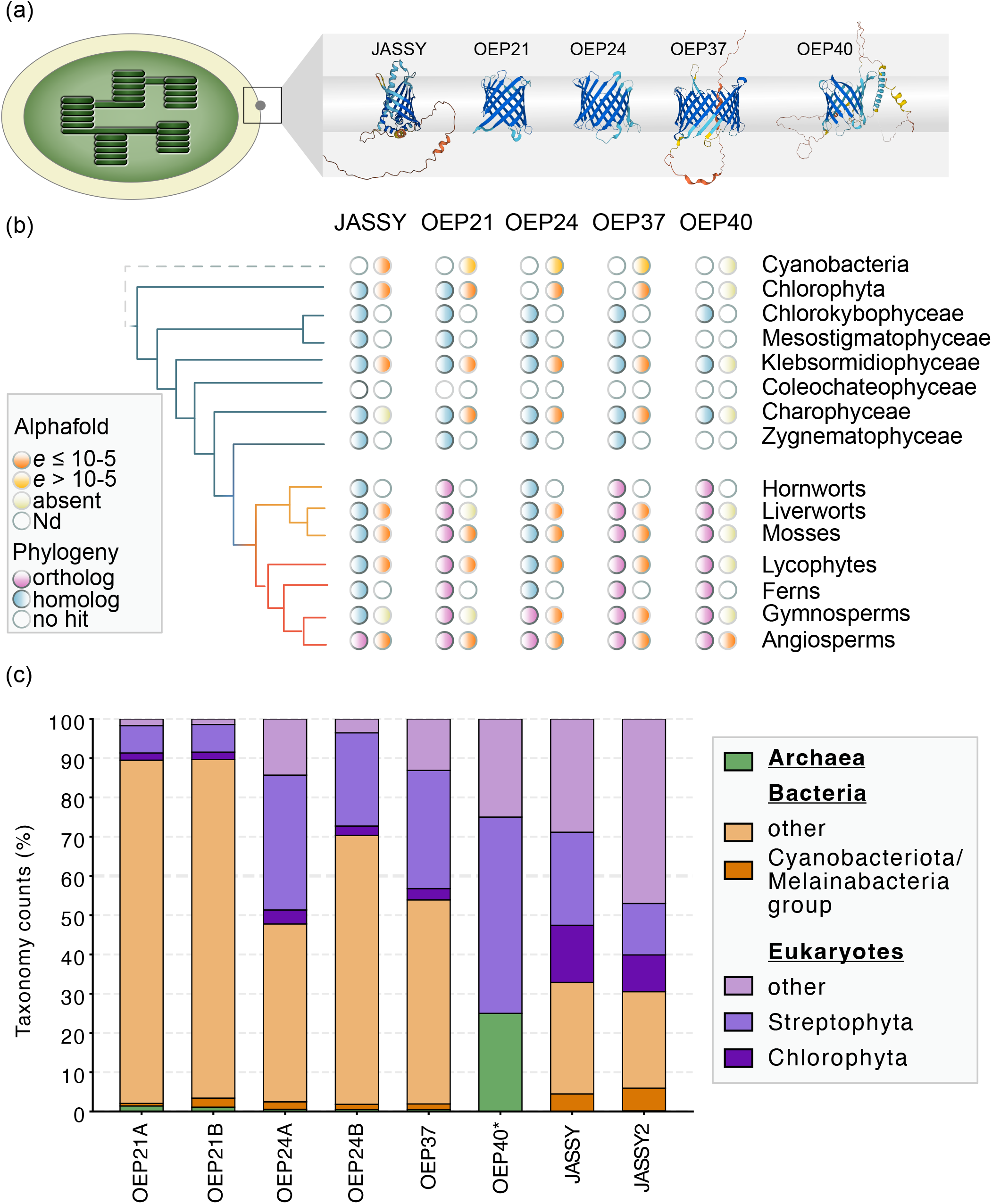
Model of key outer envelope proteins. (a) Predicted structures of proteins whose evolutionary histories were traced in this study. (b) Using primary amino acid sequence information, homologs (blue dot) and orthologs (purple dot) of JASSY, OEP21, OEP24, OEPE37, and OEP40 were detected across major chloroplastidal lineages (cyanobacteria as outgroup) using phylogenetic analysis; a white dot means that no homolog was detected. Additionally, protein structural data were used to detect structural homologs/analogs considering only hits with some structural confidence (pLDDT≥70; with the exception for OEP40; here only 5 hits were recovered and all were considered, indicated by *); to distinguish true (likely homologous) and false (likely analogous) positive hits we applied an e-value cutoff of ≤10 (orange dot). If a plant lineage only had hits with e-values >10 we annotated this with yellow dots. Absence of a structural homolog/analog among proteins with structural confidence (pLDDT≥70) is indicated by a light yellow dot (for OEP40 all hits were considered, indicated by *); a white dot indicates that no hits could be determined (Nd) because no structural data were available for the respective clade. (c) Taxonomic distribution of structural hits with a pLDDT≥70 with archaea in green, bacteria in orange and eukaryotes in purple.

To identify sequence homologs of five OE channels from Arabidopsis, we employed two approaches, BLASTp and Homology detection & structure prediction by HMM-HMM comparison (HHPred). While BLASTp uses sequence-based comparison, HHPred aims to identify homologs using sequence, domain, and in some cases structural comparison (Söding et al. 2005). HHPred identified hits with probabilities ≥70% across diverse phyla for all OEPs investigated, yet e-values were mostly high (Figure S1, Supplementary Table S1a-i). *At*JASSY1/2 had several well-supported hits to bacteria or tracheophytes, however no hits to cyanobacteria were recorded. *At*OEP21A/B had one hit with an e-value ≤ 10^−7^ to *Pisum sativum* OEP21. All other candidates retrieved only hits with insufficient support (e-values > 0.09).

To understand whether any of the OEPs can be traced to lineages other than land plants, we further conducted BLASTp searches against the entire non-redundant protein sequence (nr) database, which is more comprehensive compared to the one used in the HHPred approach. Here, we employed strategies to filter for (i) all sequences outside of Viridiplantae, (ii) rhodophytes, (iii) glaucophytes, (iv) cyanobacteria and (v) bacteria. Apart from OEP21A, which recovered possible distant homologs in the rhodophyte genus *Galdieria* (e-value ≥ 0.002 and identity <27%), none of the β-barrel proteins recovered hits in any of the categories (Supplementary Table S2a). In accordance with the HHPred approach, BLASTp also recovered one bacterial hit for JASSY, but all cyanobacterial hits had e-values ≥10^−5^ and were only recovered when specifically querying cyanobacteria (Supplementary Table S2a). However, these potential cyanobacterial sequence hits all represent START (StAR-related lipid-transfer)/RHO_alpha_C/PITP/Bet_v1/CoxG/CalC (SRPBCC) family proteins. Indeed, all cyanobacterial hits only showed similarity to the SRPBCC region of JASSY1. This suggests that they (i) could be distant homologs to JASSY from the phylogenetic twilight zone and may be better recovered using a structural approach or (ii) may not be true homologs but rather share an ubiquitously present START domain.

Apart from JASSY, which had few hits to rhodophytes and a bacterial hit, none of the other candidates seem to identify obvious sequence homologs outside of land plants. To scrutinize the appearance and sequence evolution of these OEPs, we next conducted phylogenetic analyses based on the results from BLASTp against our in-house Chloroplastida database (Figure 1, 2, S2, S3, S4, S5, Supplementary Table S2b-f). This analysis indicates that *At*OEP21A/B do not have homologous sequences beyond chlorophyte algae. Moreover, homologs to *At*OEP24A/B, *At*OEP37 and *At*OEP40 appear limited to streptophytes (Figure 1b). Clear orthologs for *At*OEP21, *At*OEP37 and *At*OEP40 exist in all embryophytes, suggesting their existence since the last common ancestor (LCA) of land plants. In contrast, OEP24 only shows clear orthology in gymno- and angiosperms, suggesting an appearance in the LCA of seed plants (Figure 1b). Taken together, our data point to an origin of the gene families in the LCAs of Chloroplastida (homologs to OEP21) or Streptophyta (homologs to OEP24, 37 and 40) and the origin of the respective subfamilies in the LCAs of land plants (OEP21, 37 and 40) and seed plants (OEP24).

**Figure 2:**
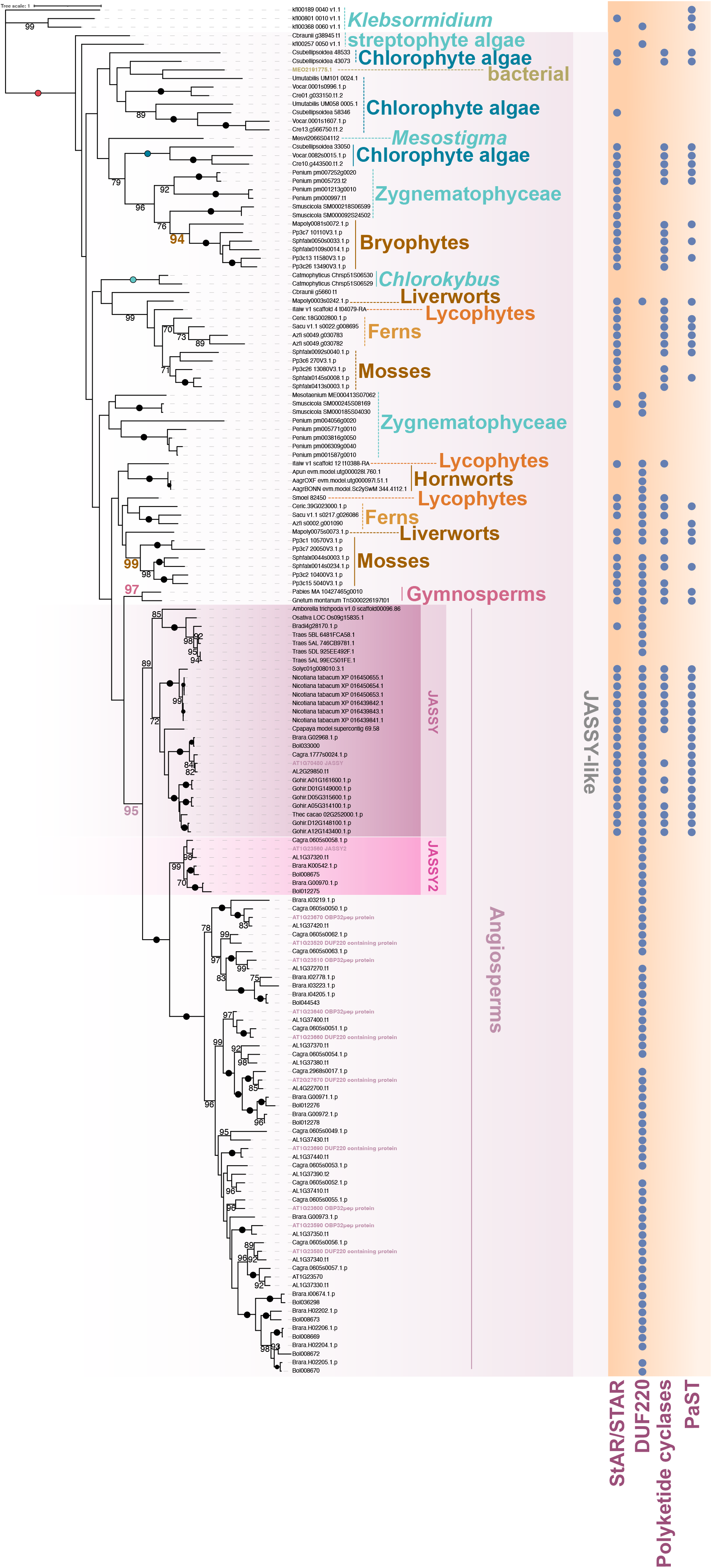
Maximum likelihood phylogeny of JASSY homologs. The phylogeny was computed based on a global primary sequence alignment using a G-INS-I approach in MAFFT. The dot matrix on the right labels which domains are predicted to be present in the protein homologs. 100 non-parametric (Felsenstein) bootstrap pseudoreplicates were computed.

Due to its unique structure and available knowledge on the evolution of jasmonate biosynthesis, we focused on JASSY1/2 in more detail (Figure 1a, Figure 2). Using BLASTp, we obtained one bacterial hit for JASSY that met our e-value threshold of <10^−7^ (Supplementary Table S2a). This hit clustered within the JASSY-like clade (bootstrap support 100) but has none of the domains recovered for JASSY-like proteins (Figure 2). This suggests that it (i) may have a different function or (ii) is a sequencing artefact. Independent of this, our phylogenetic inference points to the LCA of angiosperms having possessed a single JASSY gene, while the gene family expanded significantly in the LCA of Brassicaceae with 12 members. This is consistent with the observation that At*jassy1* loss-of-function mutants are neither completely JA deficient nor male sterile (Guan et al. 2019). Moreover, we observe lineage-specific expansion of the JASSY family in diverse branches of the green lineage (Figure 2). The *At*JASSY1 domain composition of a STAR, DUF220, Polyketide cyclases and PaST domain is only observed for homologs from land plants (Figure 2). Brassicaceae JASSY paralogs contain only the DUF220 domain, lacking evidence of the other domains. In contrast, JASSY homologs in streptophyte algae rarely possess the DUF220, but retain representative sequences for the other three domains. No streptophyte algal homolog contains a set of all four domains. This suggests stepwise domain recruitment in the JASSY family and concomitant domain loss after duplication. Stepwise domain recruitment has been previously observed in a large-scale domain acquisition analyses across Chloroplastida and was suggested to be a mechanism to contribute to functional novelty in streptophyte stress response (Dhabalia Ashok et al. 2024). How this relates to JASSY function is so far unclear, but we note that *K. nitens*, for which the presence of OPDA was reported (Chini et al. 2023, Schmidt et al. 2024) possess distantly homologous sequences that encode DUF220 or STAR domains (Figure 2). Overall, these sequence-based phylogenetic data point to a complex evolutionary history behind the origin of the chloroplast OE channels of land plants with a stepwise appearance of OEPs. Whether structurally similar cyanobacterial proteins were lost and replaced by newly appearing ones or whether sequences degraded so much that we simply cannot trace their origin back to cyanobacteria, remains unclear.

### Implementation of structural data suggest distinct evolutionary trajectories of β-barrel OEPs after stepwise recruitment

To gain a deeper understanding of the evolutionary trajectory of the OE, we employed structure similarity searches using the AFDB50 database with AlphaFold2 predictions on the selected OEPs. While β-barrel structures can be hard to predict, these challenges appear to have been overcome since AlphaFold2 (Topitsch et al. 2024). Overall, our analysis revealed that most β-barrel OEPs display a broad phylogenetic distribution based on their structure, using a predicted local distance difference test (pLDDT) cutoff ≥70 to only retain structures with confidence in their prediction (Figure 1c, Supplementary Table S3c-h). Consistent with the tree-of-life wide distribution of β-barrel structures, we found conservation across diverse bacterial, archaeal and eukaryotic sequences outside the green lineage (Figure 1c). Noteworthy, the four β-barrel OEP families did not retrieve the same taxonomic coverage or the same hits (supplementary table S3, average overlap between hits across all five OEP families: 11.3±19.3%; supplementary table S4). Thus, it appears for OEPs it is possible to distinguishing different β-barrel structures.

The only exception to the broad distribution of structurally similar sequences was *At*OEP40, which had only five structural hits. One of these had a pLDDT of 71, others were below this threshold (Supplementary Table S3h). This particular gene was annotated as a neuronal cell adhesion molecule-like protein and e-value was high (0.0062), indicating that the proteins are unlikely related. Because *At*OEP40 exhibits structural differences from *At*OEP21, 24, and 37 (Figur1a), we hypothesize that it may serve a distinct function. Additionally, we do not recover hits to (i) any of the other OEPs from Arabidopsis or (ii) an overlapping hit spectrum across eu- and prokaryotes with other OEPs on sequence and structure level (Supplementary Table S2a, S2f, S3h, S4). This further supports the lack of sequence or structural similarity with any of the other three OEPs or possible bacterial structure homologs. Altogether, our finding suggests that OEP40 may (i) have significantly diverged in structure and sequence or (ii) originated *de novo* in the last land plant common ancestor. Given that we observe sequence similarity among land plants, the absence of structural similarity to streptophytes (only two significant putative structural homologs with low pLDDT) suggests low structural conservation and rapid structural turnover (Figure 1b,c, Figure S5). Coherently, we observe low sequence similarity in the C-terminal region of OEP40 across land plants resulting in an unstructured C-terminal region (Figure 1a, S6). This points to neofunctionalization within land plants, driven by structural divergence. The function of OEP40 may thus have varying functions across land plant diversity. This makes *At*OEP40 an excellent candidate to study substrate specificity and potential evolutionary changes in this regard.

Among OEP21A/B, 24A/B, and 37, OEP24A and B demonstrated predicted structural conservation (e-value ≤10^−5^) not only within Chloroplastida but also with rhodophytes, and members from haptophytes and SAR group (Stramenopiles/Alveolates/Rhizaria) that harbor plastids via secondary endosymbiosis (Supplementary Table S3e). This result differs from the sequence-based searches (Supplementary Table S2a) and leaves us with two possibilities: we observe (i) an example of low sequence and high structural conservation or (ii) convergent folding. Haptophyte and SAR species with significant structural hits retain plastids of red algal origin (Sibbald and Archibald 2020), which may support a deep homologous ancestry and acquisition early on via EGT. Indeed, for OEP24 hits, putative functions associated with the identified hits are primarily related to porins found in organellar membranes. OEP37 has significant structural hits from all domains of life (e-value ≤10^−5^, Supplemtary Table S3g). Additionally, we found a significant structural homolog from Crysophytes (e-value ≤10^−5^) for OEP37. Crysophytes also have a red-algal origin of their plastid (Sibbald and Archibald 2020) and indeed structural hits similar to OEP37 from rhodophytes that barely missed the e-value cutoff exist. These similarities to protein structures in rhodophytes is not apparent for OEP21A/B. This is inconsistent with the low-quality hits retrieved for OEP21A from *Galdieria* via BLASTp; indicating that these BLASTp-derived hits are not deep homologs from the so-called twilight zone.

Given that the three OEP families (OEP21, 24, and 37) (i) do not recover the same taxonomic scope of structural homologs/analogs and (ii) recovered hits with distinct descriptions among the best-fitting structural models, we reason that they have distinct evolutionary histories. The analyses of these four β-barrel OEP families suggests that OEPs next to the stepwise recruitment during the evolution of the chloroplast OE, also underwent different degrees of sequence- and structural turnover. While not recovered using sequence analyses only, structural data suggest that OEP24 may be acquisitions from early EGT event. While all of the four β-barrel OEP are annotated as solute channels, their distinct functions are still largely unclear. Nevertheless, the fact that they follow different evolutionary paths, implicates that they may fulfil different functions in the chloroplast OE and that function can also shift during evolution.

### JASSY originated from latent genetic potential in the LCA of land plants

JASSY proteins exhibit unique structural configurations among OEPs: They unite β-sheets and α-helices, distinguishing them from β-barrel OEPs. Among all five OEP families, JASSY showed the broadest taxonomic distribution in the sequence-based analyses (Figure 1b, Supplementary Table S2a,b,c). JASSY1/2 encode a StAR/START domain, which is typical for lipid transporters, but also other lipid binding proteins, likely causing the broad taxonomic scope. Both HHPred and Alphafold results support this assumption. This is coherent with a recent comparison between sequence and structural methods showing that Foldseek-based structural searches tend to identify a high number of non-homologous sequences (Mutti et al. 2024). Highly conserved domain sets may be a reason for that.

To scrutinize the evolutionary history of JASSY1 and JASSY2, we worked with the structural models of Alphafold2 and computed phylogenies based on structure-based alignments (Figure 3). We recovered three distinct clades with broader streptophyte representation, one clade containing homologs to JASSY and the other two clades containing homologs to *At*PPP1 and a polyketide cyclase, supporting that several initial hits were attracted due to common domains. Cyanobacterial sequences did not cluster with the JASSY homologs, suggesting that the BLASTp hits discussed above are not distant homologs from the twilight zone, but rather other lipid-binding proteins. Bacterial hits that cluster with the chlorophytes in both phylogenetic approaches likely signify a HGT from eukaryotes to bacteria. Consistent with the BLASTp results we identified significant structural hits from rhodophytes that fell into the JASSY-like clade (Figure 3a) Here, too, we found homologous hits from haptophytes and ochrophytes, suggesting that JASSY-like homologs were already present prior to the split of Rhodophyta and Chlorophyta. Moreover, we identified structural hits from Lokiarcheota opening the possibility for an ancient origin. The chlorophyte and streptophyte representation of the JASSY- and JASSY-like containing clade was highly similar to that in the sequence-based phylogeny adding confidence to protein-structure based approaches, even though these approaches are still in development (Garg and Hochberg 2024). In our report, the structural phylogenies were successful in aiding to understand uncertainties and provide further evolutionary insight by combining the use of structure as a baseline for sequence alignments and only sequence-based approaches.

**Figure 3:**
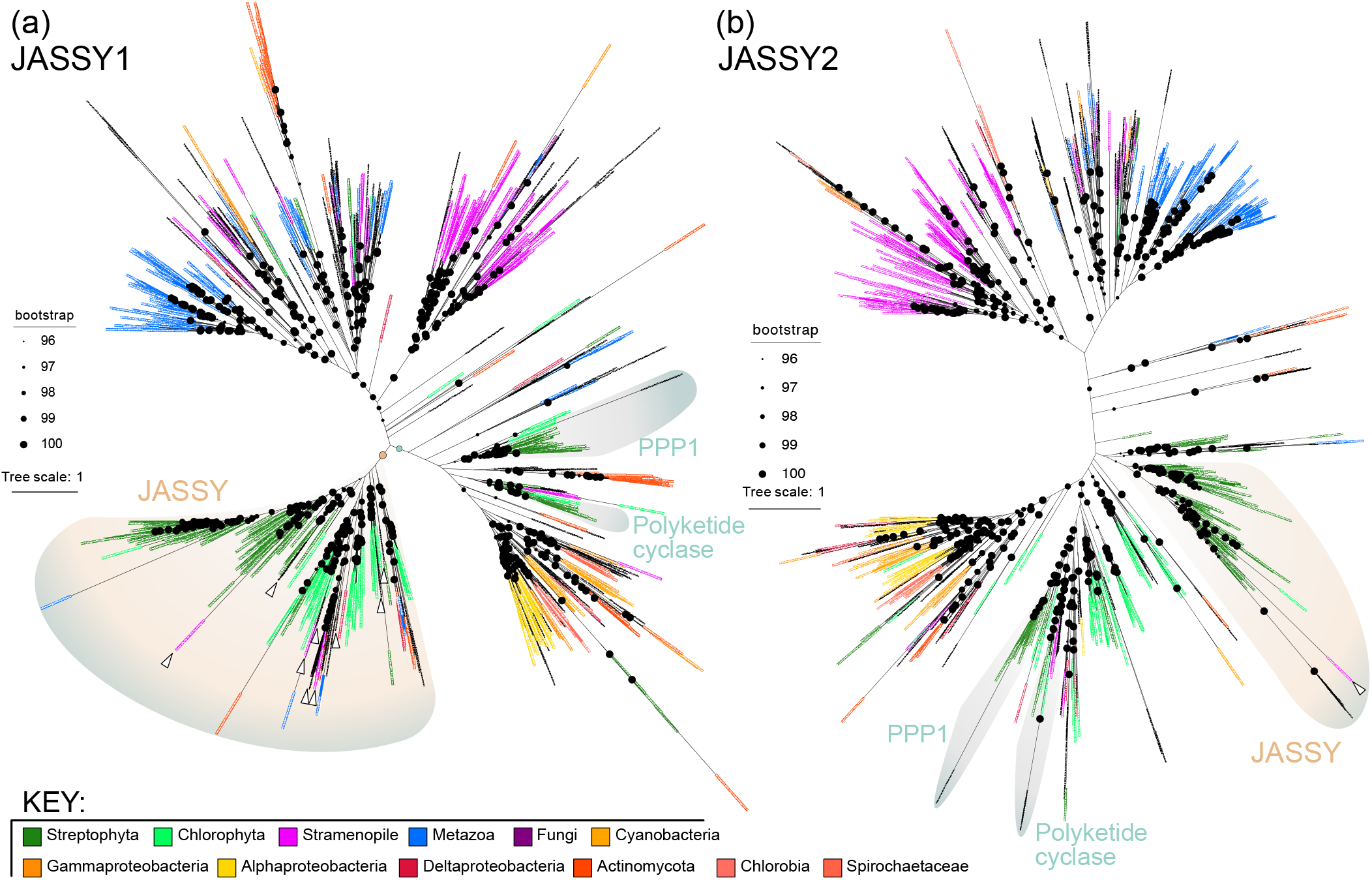
Composite phylogenetic analysis of JASSY homologs based on primary sequence and structural protein data. Maximum likelihood phylogenies of structure and sequence (a) JASSY1 and (b) JASSY2 combining information derived from amino acid distance matrices and 3Di (for details see M&M). Structural information is based on AlphaFold database search. 1000 ultrafast bootstrap pseudoreplicates were computed. White arrows indicate homologs from rhodophytes and species with a secondary endosymbiosis event.

The only characterized member of the JASSY family is *Arabidopsis thaliana* JASSY1 facilitating OPDA export from chloroplasts (Guan et al. 2019). But when did this function of JASSY evolve? Our domain analysis across JASSY homologs suggests that the JASSY1 domain setup was first recruited in the LCA of land plants, while JASSY-like sequences predate this occurrence (Figure 2). The similarity to other StAR/START and DUF220-containing proteins in structural or domain-based analyses supports an ancient origin of the domains followed by lineage-specific domain duplications and distributions across the genomes. Yet, despite the absence of the StAR/START domain our structural phylogeny supported a cluster of JASSY-like proteins from chlorophytes and streptophytes.

Among the structurally similar hits, hits to putative transporters of non-polar substrates have been annotated in other bacteria and eukaryotes within the AFDB50 database. To our knowledge, screens across the green lineage have detected neither OPDA nor dn-OPDA in chlorophytes (Chini et al. 2023, Schmidt et al. 2024) and distribution of them in streptophyte algae was scattered (Schmidt et al. 2024). Further, the overlap between the algal sequences we included in our analyses and the species analyzed for OPDA and dn-OPDA production is small. More metabolomic screenings of chlorophyte and streptophyte algae with JASSY homologs and functional characterization of these deep homologs are needed. Without these, an inference of the functional necessity of any of the four domains that bona fide JASSY proteins encode remain obscure. Regarding the evolutionary history of the JASSY-like family, we hypothesize that JASSY emerged from a novel domain assembly. It is likely that the progenitor of JASSY was a transporter for long-chain, non-polar substances that was co-opted in the green lineage to become an OPDA transporter in land plants.

## Conclusion

OEPs are key elements for chloroplast function. However, tracing their evolutionary history is challenging: For one, the cyanobacterial progenitor of plastids was a genetic chimera shaped by genes acquired through HGT. This complicates efforts to definitively trace genes to their cyanobacterial ancestry, even when that is their true origin (Ku et al., 2015). Second, OEPs may have been co-opted from host-derived proteins. Here, we combined structure-based phylogenetic approaches with traditional phylogenetic analyses to understand the evolutionary history of five OEP families. All OEPs, except OEP40, have clear structural similarities to sequences found across the tree of life. However, except for JASSY, none of this is reflected on the sequence level. The OEPs analyzed here likely resulted (i) from neofunctionalization of pre-existing protein families after ancient duplication events or domain reshuffling, the latter being likely the case for JASSY and/or (ii) ancient EGTs—a possibility for OEP24 family. The presence of structurally similar sequences across diverse life forms, coupled with the difficulty in tracing most OEPs’ sequence-level origins, highlights the intricate evolutionary pathways these proteins have traversed. It also emphasizes the power of combined structure and sequence-level analyses in unraveling the evolutionary history of these crucial chloroplast proteins. Our analyses only scratch the surface of the evolutionary space and structure of OEPs or other plastid transporter proteins. Yet, it is clear from our data that OEPs did not originate all at the same time: Some have been recruited later than others, were co-opted or may have replaced existing bacterial versions due to structural similarities. We now know that the evolution of the OE protein make-up is dynamic, although the driving forces of these dynamics remain obscure. Going forward, we need to integrate structural and sequence-based approaches with biochemical and physiological data to study these evolutionary dynamics that gave rise to today’s chloroplast.

### Material and Methods Sequence homology analyses

We used the *A. thaliana* protein sequences from JASSY1 and 2, OEP21A, B, OEP24A, B, OEP37, and OEP40 to query our database of land plants and streptophyte and chlorophyte algae (Organisms included: (*Anthoceros agrestis* Oxford and Bonn (Li *et al*., 2020), *Anthoceros punctatus* (Li *et al*., 2020), *Amborella trichopoda* (Amborella Genome Project *et al*., 2013), *A. thaliana* (Lamesch *et al*., 2011), *A. lyrata* (Hu *et al*., 2011), *Azolla filiculoides* (Li *et al*., 2018), *Bathycoccus prasinos* (Moreau *et al*., 2012), *Brassica oleracea* (Liu *et al*., 2014), *Brassica rapa* (Goodstein *et al*., 2011), *Brachypodium distachyon* (Vogel *et al*., 2010), *Capsella grandiflora* (Slotte *et al*., 2013), *Carica papaya* (Ming *et al*., 2008), *Ceratopteris richardii* (Marchant *et al*., 2022), *Chara braunii* (Nishiyama *et al*., 2018), *Chlorokybus melkonianii* ((Wang *et al*., 2020); for naming, see (Irisarri *et al*., 2021)), *Chlamydomonas reinhardtii* (Merchant *et al*., 2007), *Coccomyxa subellipsoidea* (Blanc *et al*., 2012), *Gnetum montanum* (Wan *et al*., 2018), *Gossypium hirsutum* (Li *et al*., 2015), *Isoetes taiwaniensis* (Wickell *et al*., 2021), *Klebsormidium nitens* (Hori *et al*., 2014), *Marchantia polymorpha* (Bowman *et al*., 2017), *Mesostigma viride* (Wang *et al*., 2020), *Micromonas pusilla* (Worden *et al*., 2009), *Micromonas* sp. (Worden *et al*., 2009), *Nicotiana tabacum* (Sierro *et al*., 2014), *Oryza sativa* (Ouyang *et al*., 2006), *Picea abies* (Nystedt *et al*., 2013), *Physcomitrium patens* (Lang *et al*., 2018), *Salvinia cucullata* (Li *et al*., 2018), *Selaginella moellendorffii* (Banks *et al*., 2011) *Solanum lycopersicum* (Sato *et al*., 2012), *Sphagnum fallax* (Healey *et al*., 2023), *Theobroma cacao* (Argout *et al*., 2011), *Triticum aestivum* (The International Wheat Genome Sequencing Consortium *et al*., 2018), *Mesotaenium endlicherianum* V1 (Cheng *et al*., 2019), *Ostreococcus lucimarinus* (Palenik *et al*., 2007), *Penium margaritaceum* (Jiao *et al*., 2020), *Spirogloea muscicola* (Cheng *et al*., 2019), *Ulva mutabilis* (De Clerck *et al*., 2018), *Volvox carteri* (Prochnik *et al*., 2010)) using BLASTp. We filtered the output according to e-value and only retained hits with an e-value of ≥10. Additionally, we queried the NCBI nr database with the following restrictions (i) excluding Viridiplantae, (ii) excluding Viridiplantae and bacteria, (iii) Cyanobacteria, (iv) Rhodophyta and (v) Glaucophyta.

We used protein sequences of JASSY1 and 2, OEP21A, B, OEP24A, B, OEP37, and OEP40 as an input in HHPred (Zimmermann et al. 2018, Gabler et al. 2020). The search was made across the default database PDB_mmCIF70_16_Aug for each protein separately. No specific proteomes were selected. We used the default parameters in the search except E-value cutoff, which changed to 0.05 and the number of max target hits increased to 10000. After downloading the results, we created a table with organism names and the probabilities for the hits. Then these tables were used to plot histograms in figure S1. To check if cyanobacteria hits are present or not, we used a script to find phylum of each species (https://github.com/derenbuyuktas/Find_Phylum).

### Phylogenetic reconstruction based on sequence similarity approaches

We fused the output of the BLASTp analyses and aligned all sequences belonging to the same family (in total five families: JASSY, OEP21, OEP24, OEP37, and OEP40) using mafft with a G-INS-I approach (Katoh and Standley 2013). We checked the quality of the alignment and proceeded to calculating a ML phylogeny. For phylogenetic reconstruction of the BLASTp results, we used IQ-TREE with 100 bootstraps per phylogeny (Nguyen et al. 2014). Models were selected using modelfinder implemented in IQ-TREE (Kalyaanamoorthy et al. 2017). Sequence-based phylogenies were visualized with iTOL (Letunic and Bork 2021) and annotated manually for taxonomy.

### Structural predictions

We used Foldseek (van Kempen et al. 2024) to identify structurally similar proteins using the Alphafold database (AFDB50, Jumper et al. 2021, Varadi et al. 2024, accessed 29 October 2024). As query we used the protein sequences of *A. thaliana* for JASSY1 and 2, OEP21A, B, OEP24A, B, OEP37, and OEP40. For those were several models existed we used the one with the highest quality (UniProt golden star or if not applicable the highest sequence similarity across most of the protein to the variations that existed for the *Arabidopsis* proteins in the database). We next selected a cutoff of pLDDT≥70 to include high-confidence structural models, while allowing disordered side-chains. In downstream analyses we further applied an e-value cutoff of 10 to distinguish likely homologous structural hits from analogous structural hits. Taxonomic identification of hits was done by recovering the TaxID for the genus a particular hit corresponded to.

### Domain predictions

Domains were predicted using Interpro and CD batch search using the Arabidopsis thaliana protein sequences as query. Both tools gave the similar results. Significant domains were only reported for JASSY1 and 2. OEP21A/B, 24A/B, and OEP37 had domains named after them, spanning the whole protein, e.g. OEP21A and 21B encode the domain OEP21. Such domains were not visualized. OEP40 included the OS08G0510800 PROTEIN, which was not identified using CD search.

### Phylogenetic reconstruction based on structural alignments

We, here, follow a novel approach for structural phylogenetics of JASSY1 and 2 first presented in Puente-Lelievre et. al (2024), which utilizes the 3Di alphabet introduced by Foldseek to capture tertiary interactions between neighboring residues as information for phylogenetic reconstruction in addition to standard amino acid alignments using a partition scheme in a maximum likelihood framework for a detailed discussion see Puente Lelievre et. al (2024) and Garg & Hochberg (2024).

To infer the structural phylogeny for JASSY1 and 2, we used the the list of possible homologs of JASSY1 and 2 derived from the search in the Alphafold database, we selected hits with an e-value ≥10. The putative homologs were aligned with the structural aligner FoldMason (Gilchchrist et al. 2024) to get both the amino acid and 3Di alignments. The alignments were then trimmed using ClipKIT -m gappy (Steenwyk et al. 2020) in the case of amino acid alignments and trimAl (Capella-Gutiérrez et al. 2009) -gappyout in the case of 3Di alignments, where trimAl performs more extensive trimming in order to confine the 3Di analysis to the protein cores. Trees were constructed using IQ-TREE 2.4.0 and using Modelfinder to find the best fit substitution model and 1000 Ultrafast bootstraps (Hoang et al. 2018). For the combined amino acid and 3Di analysis, both alignments were concatenated and run with an edge-proportional partition scheme (Chernomor et al. 2016), where the custom substitution matrix by Garg and Hochberg (2024) is specified for the 3Di alignment.

## Acknowledgement

We thank Johannes Söding (MPI for Multidisciplinary Sciences, Göttingen) for constructive feedback regarding the setup for structural analyses. SdV is grateful to the German Research Foundation (DFG) for funding (Cymbiomics, VR139/2-1; 515101361). SS received funding from the DFG (CRC TR175, project B06).

