## Supplementary material for "The evolutionary history of plastid outer envelope proteins – a structure-sequence comparison": Figure S1

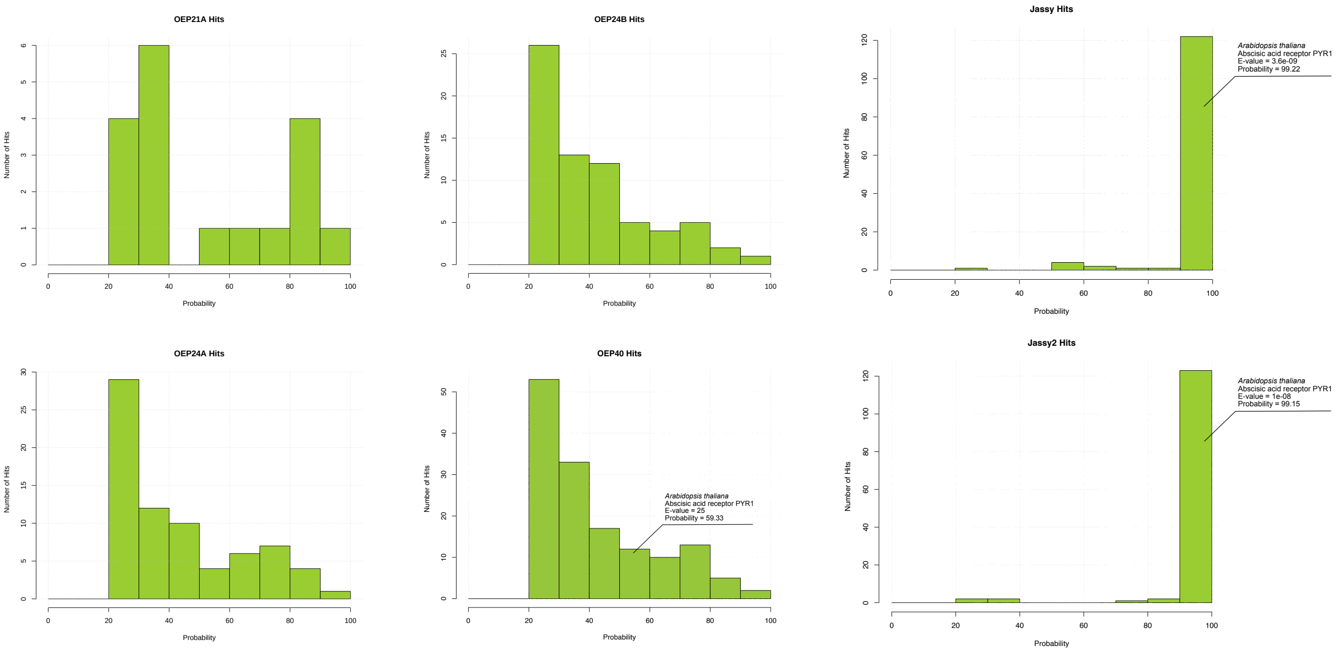


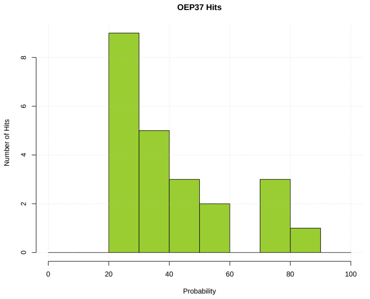


**Figure S1. HHPred probability distribution for hits of putative OEPs across the tree of life.** With exception of JASSY1 and 2, the other OEPs recovered only low-probability hits. The best hit to a protein from *Arabidopsis thaliana* other than the query is indicated. No cyanobacterial hits were found for any of the 7 queried proteins. The next best hit in *Arabidopsis thaliana* was, if one occurred, *At*PYR1, although probability was low for it in case of OEP40. It nevertheless indicates high domain similarity to some of the query proteins. The probability bins are given on the x-axis and the y-axis represents the number of sequences within a particular probability bin.
