## Supplementary material for "The evolutionary history of plastid outer envelope proteins – a structure-sequence comparison": Figure S3

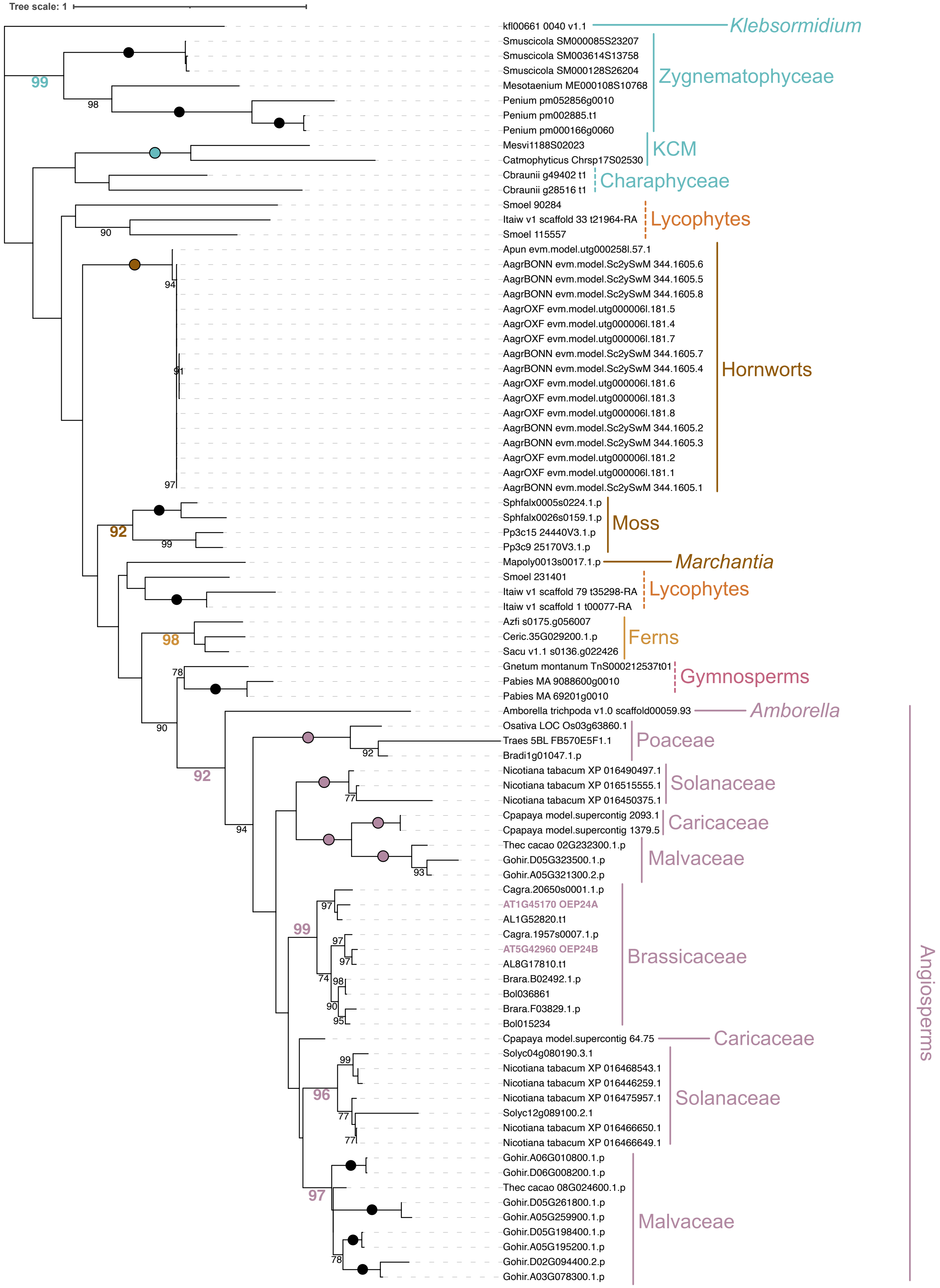


**Figure S3. Phylogenetic analyses of OEP24.** The unrooted phylogeny is based on BLASTp hits of *At*OEP24 and B against a streptophyte and chlorophyte genomes containing database and the nr database on ncbi restricted to (a) bacteria and (b) cyanobacteria. *Arabidopsis* gene family members OEP24A and OEP24B is highlighted in light purple and bold. Taxonomic affiliation is indicated on the right. Bold lines show monophyletic groups with a minimum bootstrap support of >75. Dotted lines indicate sequences belonging to the same lineage, which are either paraphyletic or not supported to form a monophylum. Only bootstrap support of >75 is shown, bootstrap support of 100 is indicated by circles.
