## Supplementary material for "The evolutionary history of plastid outer envelope proteins – a structure-sequence comparison": Figure S6

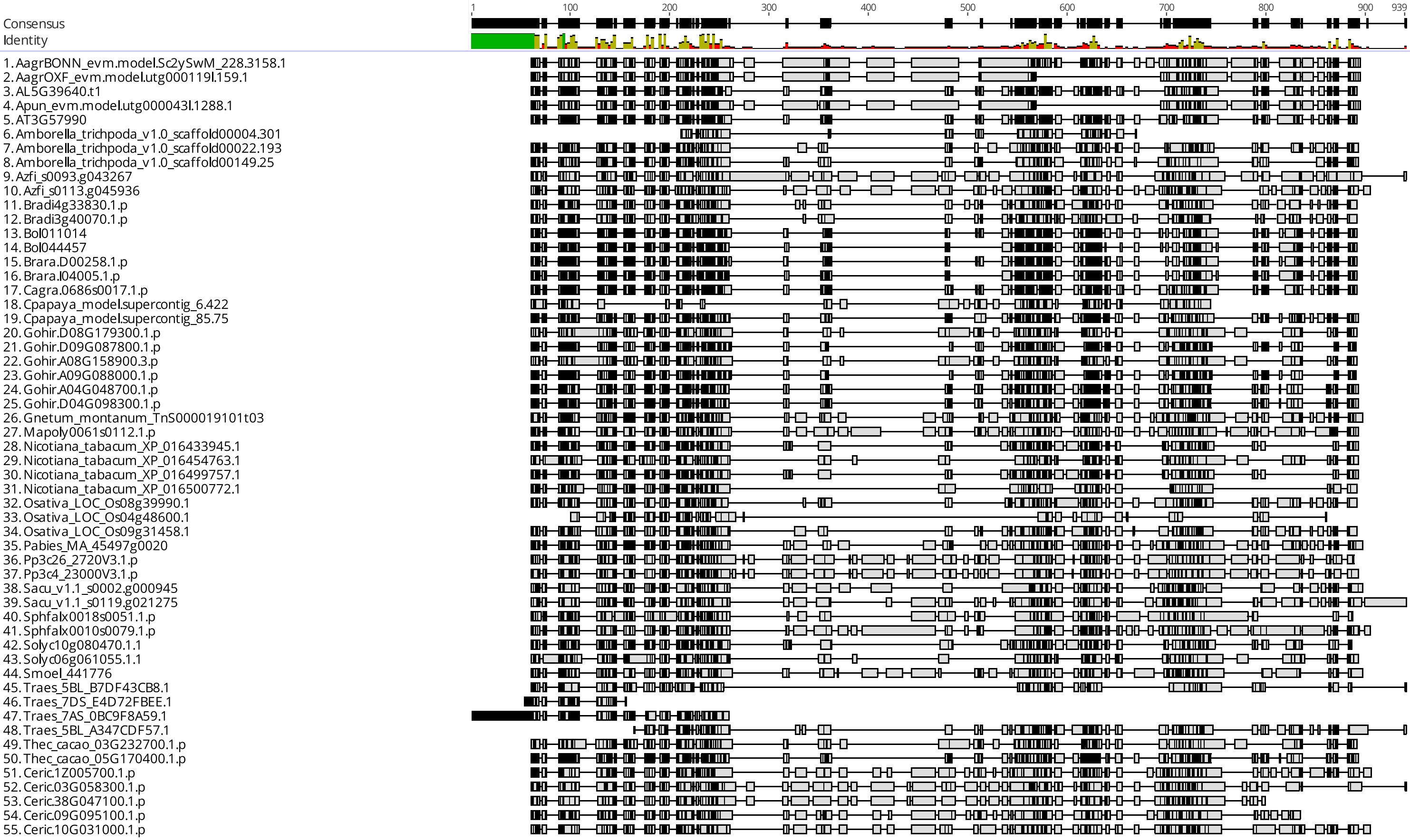


**Figure S6. Alignment of OEP40 homologs from land plants.** On top consensus sequence and identity histogram with identity score coloring from red (low) to green (high) across all amino acids at a given position is visualized. Gaps are not considered for the identity score. Below is a schematic overview of the G-INS-I alignment of OEP40 homologs. Residues in black show conserved residues, residues in grey are not conserved in relation to the consensus sequence.
